# Formaldehyde-responsive proteins, TtmR and EfgA, reveal a tradeoff between formaldehyde resistance and efficient transition to methylotrophy in *Methylorubrum extorquens*

**DOI:** 10.1101/2020.10.19.346494

**Authors:** Jannell V. Bazurto, Eric L. Bruger, Jessica A. Lee, Leah B. Lambert, Christopher J. Marx

## Abstract

For bacteria to thrive they must be well-adapted to their environmental niche, which may involve specialized metabolism, timely adaptation to shifting environments, and/or the ability to mitigate numerous stressors. These attributes are highly dependent on cellular machinery that can sense both the external and intracellular environment. *Methylorubrum extorquens* is an extensively studied facultative methylotroph, an organism that can use single-carbon compounds as their sole source of carbon and energy. In methylotrophic metabolism, carbon flows through formaldehyde as a central metabolite; thus, formaldehyde is both an obligate metabolite and a metabolic stressor. Via the one-carbon dissimilation pathway, free formaldehyde is rapidly incorporated by formaldehyde activating enzyme (Fae), which is constitutively expressed at high levels. In the presence of elevated formaldehyde levels, a recently identified formaldehyde-sensing protein, EfgA, induces growth arrest. Herein, we describe TtmR, a formaldehyde-responsive transcription factor that, like EfgA, modulates formaldehyde resistance. TtmR is a member of the MarR family of transcription factors and impacts the expression of 75 genes distributed throughout the genome, many of which are transcription factors and/or involved in stress response, including *efgA.* Notably, when *M. extorquens* is adapting its metabolic network during the transition to methylotrophy, *efgA* and *ttmR* mutants experience an imbalance in formaldehyde production and a notable growth delay. Although methylotrophy necessitates that *M. extorquens* maintain a relatively high level of formaldehyde tolerance, this work reveals a tradeoff between formaldehyde resistance and the efficient transition to methylotrophic growth and suggests that TtmR and EfgA play a pivotal role in maintaining this balance.

**Importance:** All organisms produce formaldehyde as a byproduct of enzymatic reactions and as a degradation product of metabolites. The ubiquity of formaldehyde in cellular biology suggests all organisms have evolved mechanisms of mitigating formaldehyde toxicity. However, formaldehyde-sensing is poorly described and prevention of formaldehyde-induced damage is primarily understood in the context of detoxification. Here we use an organism that is regularly exposed to elevated intracellular formaldehyde concentrations through high-flux one-carbon utilization pathways to gain insight into the role of formaldehyde-responsive proteins that modulate formaldehyde resistance. Using a combination of genetic and transcriptomic analyses, we identify dozens of genes putatively involved in formaldehyde resistance, determined the relationship between two different formaldehyde response systems and identified an inherent tradeoff between formaldehyde resistance and optimal transition to methylotrophic metabolism.

## INTRODUCTION

Methylotrophy, a trait found in all domains of life, is the unique metabolic ability of organisms to use one- or multi-carbon compounds lacking carbon-carbon bonds, as sole sources of carbon and energy. Growth substrates for methylotrophs include compounds such as methane (methanotrophs), methanol, and methylamines (mono-, di-, tri-), and methylated sulfur species. A key feature of methylotrophic metabolism is that each carbon of the growth substrate flows through the potent toxin formaldehyde as a central intermediate. Formaldehyde is harmful to all cell types because of its reactivity with electrophiles such as free amines and thiols. It can form adducts with or crosslink a number of biological molecules such as DNA and proteins. Despite the centrality of formaldehyde in methylotrophic metabolism, until very recently formaldehyde-specific stress responses had not been described in methylotrophs.

In *Methylorubrum* (formerly *Methylobacterium) extorquens* AM1 the most extensively studied facultative methylotroph, the intracellular formaldehyde concentration is estimated at ~ 1 mM during growth on methanol (1). Methanol is directly oxidized to formaldehyde in the periplasm by methanol dehydrogenase (MDH). Free endogenous formaldehyde is kept relatively low in the cytoplasm by formaldehyde activating enzyme (Fae), which condenses formaldehyde with the C1 carrier dephosphotetrahydromethanopterin (dH4MPT); this is the first of three reactions in the pterin-dependent dissimilatory pathway that oxidizes formaldehyde to formate (2–4). Previous work has demonstrated that strains defective in formaldehyde dissimilation by disruption of pathway enzymes (*fae*) or synthesis of the dH4MPT C1 carrier (*mptG*) are sensitive to methanol (4). Formate is the branchpoint metabolite and can be further oxidized to CO2 for energy production or shunted toward a tetrahydrofolate-dependent assimilation pathway for ultimate incorporation into biomass via the serine cycle (5, 6).

Very recently experimental evolution of *M. extorquens* PA1 for growth on formaldehyde as a sole source of carbon and energy revealed multiple loci that could increase formaldehyde resistance (7). Although dissimilation by the dH4MPT pathway is the primary mechanism for removing free formaldehyde from the cytoplasm (3, 4), none of the beneficial mutations were in these genes. Instead, this work found that single mutations in one of three genetic loci *(efgA, def, Mext_0925)* could independently confer the ability to use formaldehyde as a sole source of carbon and energy. The most frequent class of mutations (nearly 80%) were loss-of-function mutations in **<u>e</u>**nhanced **<u>f</u>**ormaldehyde **<u>g</u>**rowth protein **<u>A</u>** (EfgA), a protein that directly binds formaldehyde and halts translation in the presence of excess formaldehyde by an unknown mechanism. Though strains lacking EfgA can grow in the presence of higher formaldehyde concentrations than wild-type, the absence of EfgA increases methanol sensitivity in strains that are vulnerable to formaldehyde stress (*fae*, *mptG*), suggesting that the physiological role of EfgA is to decrease formaldehyde-induced stress. Peptide deformylase (PDF, encoded by *def)*, is a ribosomally-associated enzyme that is essential for translation and involved in protein quality control (8). Selection for mutations in *def,* provided further evidence for the importance of translational regulation in response to formaldehyde stress.

Mext_0925 is a homolog of the MarR transcription factor (7). This family of regulators is found in bacteria and archaea and was originally characterized in *Escherichia coli* (9–12). In *E. coli*, MarR is encoded by the <u>m</u>ultiple <u>a</u>ntibiotic <u>r</u>esistance *(mar)* operon. It is a ligand-sensing repressor that binds the operator of the *marRAB* operon in the absence of ligand and is thus self-regulating (13, 14). A number of structurally distinct compounds such as tetracycline, chloramphenicol, and salicylate can bind MarR and induce conformational changes that lead to DNA release (19). Additionally, the *marRAB* operon can be activated by aromatic amino acid metabolites directly and indirectly (15). Subsequently, *marRAB* expression gives rise to the transcription factor MarA, a member of the AraC/XylS family, which in turn upregulates dozens of genes that contribute to resistance to multiple antibiotics, oxidative stress, and organic solvent stress (16–18). Broader examination of MarR homologs in a variety of organisms show that homologs i) can be repressors, activators, or both, ii) have highly variable regulon sizes, and iii) control a variety of cellular processes involved in various stress responses and metabolic pathways (19).

In *Bacillus subtilis*, the MarR homolog HxlR has a role in formaldehyde stress response (20, 21). *hxlR* is divergently transcribed from the *hxlAB* operon that HxlR positively regulates. The *hxlAB* operon encodes two key proteins of the ribulose monophosphate pathway of formaldehyde detoxification and is induced by exogenous formaldehyde (20, 22). *In vitro*, formaldehyde does not impact binding of HxlR and, as is the case with several MarR homologs, the mechanism of activation has remained elusive. In some instances, however, the mechanism of activation of MarR homologs is clear. For example, in organisms such as *Acinetobacter baylyi* and *Streptomyces coelicolor*, where MarR homologs regulate operons encoding enzymes required for the catabolism of aromatic lignin-derivatives such as ferulate and protocatechuate, respectively, ligand binding leads to release of cognate DNA and the operon is derepressed (19, 23–25). This leads to expression of the relevant enzymes and the ligand/carbon substrates are utilized.

Herein, we probe the physiological role of Mext_0925, implicated in modulating formaldehyde resistance. Unlike many described MarR homologs, *Mext_0925* is not in an operon or divergently transcribed from an operon, thus there is no genomic context for discerning its regulon. In the absence of obvious genetic associations, we opted to use transcriptomic analyses to identify differentially expressed genes in a *ΔMext_0925* mutant. By coupling this approach with *in vivo* genetic studies, we find that Mext_0925 is specifically responsive to formaldehyde stress. Our results define the relationship between Mext_0925 and EfgA, the only characterized formaldehyde stress response system in *M. extorquens*, and demonstrate that the ability of *M. extorquens* to respond to formaldehyde stress at the transcriptional and translational levels is critical for the optimal transition to methylotrophy. Based on our findings, we conclude that Mext_0925 represents a second formaldehyde-specific response system in *M. extorquens* and suggest the name TtmR, for a regulator of the **<u>t</u>**ransition **<u>t</u>**o **<u>m</u>**ethylotrophy.

## MATERIALS AND METHODS

### Bacterial strains, media, and chemicals

Bacterial strains used in this study (Table S1) are derivatives of *Methylorubrum extorquens* PA1 (formerly *Methylobacterium)* (26–28) where genes for cellulose synthesis were deleted to prevent aggregation and optimize liquid growth measurements (29). Therefore, the genotype referred to herein as ‘wild-type’ (CM2730) is more accurately *ΔcelABC.* The *ΔefgA* mutant (CM3745) additionally has a markerless deletion that eliminated 404 bp from the ORF of *Mext_4158* (21-424/435 bp) (7) and the *ΔttmR* mutant (CM4732) has a markerless deletion of the entire coding region (588 bp) of*Mext_0925.* All growth experiments with liquid medium were performed with *Methylobacterium* PIPES (MP) medium (29) with 3.5 mM succinate, 15 mM methanol, or 2, 4, 6, 8, 10 mM formaldehyde as a carbon source. For growth on solid MP medium, Bacto Agar (15 g/L, BD Diagnostics) was added and carbon source concentrations were elevated (15 mM succinate, 125 mM methanol). Formaldehyde stock solutions (1 M) were prepared by boiling a mixture of 0.3 g paraformaldehyde and 10 mL of 0.05 N NaOH in a sealed tube for 20 min; stock solutions were kept at room temperature and made fresh weekly. When present in the media, kanamycin was used at a final concentration of 50 μg/mL.

### Genetic approaches

Markerless deletions were generated by allelic exchange as previously described using either pCM433 (30) or pPS04_(7). Vectors were designed using SnapGene software. The HiFi assembly kit from New England Biolabs was used to construct vectors from linearized vector backbone (restriction enzyme-digested) and PCR-generated inserts.

### Growth quantitation

To initiate liquid growth, individual colonies were used to inoculate 2 mL MP medium containing 3.5 mM succinate or 15 mM methanol in biological triplicate. Cultures were shaken (250 rpm) during incubation at 30 °C for 24 hr (succinate) or 36 hr (methanol) and then subcultured (1/64) into 5 mL of identical medium for further acclimation. After this second 24-36 hr (succinate) or 36-48 hr (methanol) incubation, the stationary phase acclimation cultures were again subcultured (1/64) into relevant media for growth measurements. Cell density was determined by monitoring absorbance with a Spectronic 200 (Thermo Scientific) or a SmartSpec Plus (Bio-Rad) at 600 nm. Final yield is defined as the absorbance reached upon entry into stationary phase. Cell viability was determined by harvesting cells from a 100 μL aliquot of culture by centrifugation, discarding supernatant and resuspending the cell pellet into an equal volume of MP medium (no carbon). Cell suspensions were then serially diluted (1/10 dilutions, 200 μL total volume) in 96 well polystyrene plates with MP medium (no carbon) and 10 μL aliquots of each dilution were spotted to MP medium plates (15 mM succinate) using technical triplicates. Plates were inverted and incubated at 30 °C until colony formation was apparent (4-6 d) at which point colonies were counted. Technical triplicates were averaged for each sample; biological replicates were averaged.

### Formaldehyde quantification

Formaldehyde concentrations in the culture media were measured as previously described (31). Supernatant from a 100 μL aliquot of culture was isolated by centrifugation (14,000 x g). In technical triplicate, 10 μL of the supernatant or 100 μL of 0.1X supernatant (diluted with MP medium, no carbon) was combined with 190 or 100 μL Nash reagent B (2 M ammonium acetate, 50 mM glacial acetic acid, 20 mM acetylacetone), respectively, in 96 well polystyrene plates. Reaction plates were incubated (60 °C, 10 min), cooled to room temp (5 min), and absorbance was read at 432 nm on a Wallac 1420 VICTOR Multilabel reader (Perkin Elmer). Formaldehyde standards were prepared daily from 1 M formaldehyde stock solutions and a standard curve was read alongside all sample measurements.

### RNA-Sequencing analysis

The WT (CM2730) and the *ΔttmR* mutant (CM4732) were grown in biological triplicate in MP medium with 15 mM methanol as described above. The final growth vessel was a 250 mL flask, with a culture volume of 100 mL. When OD600 reached 0.2-0.3, cells from 50 mL of culture were harvested by centrifugation in an Eppendorf Centrifuge 5810R (5 min, 4,000xg, 4 °C). The supernatant was decanted and cells were washed with ice cold MP medium (no carbon) and centrifuged once more to remove wash supernatant. Pellets were immediately frozen by submerging tubes in liquid nitrogen and stored at −80 °C.

Nucleic acid extraction and molecular manipulation (RNA extraction, cDNA generation, library preparation) and sequencing were conducted by Genewiz (South Plainfield, NJ). The total RNA was extracted with the RNeasy Plus Mini Kit (Qiagen); rRNA was depleted with the Ribo-Zero rRNA removal kit (Illumina); the quality of resulting RNA samples was determined with an Agilent 2100 BioAnalyzer and Qubit assay. Once the cDNA library was generated, it was sequenced on an Illumina HiSeq (2×150 bp). The raw data (FASTQ format) was provided to investigators for further analysis.

An analysis pipeline developed by the University of Minnesota Genomics Center and the Research Informatics Solutions (RIS) group at the University of Minnesota Supercomputing Institute was used for data analysis. The pipeline uses FastQC (32) to assess the quality of the sequencing data, Trimmomatic (33) to remove the low quality bases and adapter sequences, HISAT2 (34) to align the curated reads to the *M. extorquens* PA1 genome [GenBank accession: CP000908.1], Cuffquant and Cufnorm from the Cufflinks package (35) to generate FPKM expression values, and featureCounts from the Rsubread R package (36) to generate raw read counts. DESeq2 was used to convert raw count data to normalized counts and all further statistical analyses were carried out in R (37). Differentially expressed genes in the *ΔttmR* mutant were identified by a pair-wise comparison to the WT strain and had a log_2_ fold change > 1 and a False Discovery Rate (FDR) adjusted p-value < 0.05.

### Stress tests

For all tests, cultures of WT (CM2730) and the *ttmR^EVO^* mutant (CM3919) were grown in biological triplicate in MP medium (succinate), as described above.

#### Alternative aldehyde growth assays

Growth was quantified in liquid media as described above with succinate or methanol as the primary carbon source. Wild-type (CM2730) was subjected to variable glyoxal, acetaldehyde, glutaraldehyde, butyraldehyde, and propionaldehyde (up to 10 mM) to identify a concentration that would lead to a growth defect but allow growth within a 24 hr period. Once the working concentrations were identified (1.25 mM for acetaldehyde and glyoxal, 2.5 mM for the remaining aldehydes), the *ttmR^EVO^* (CM3919) and *ΔefgA* (CM3745) mutants were grown in identical conditions, alongside the wild-type control.

#### Antibiotic resistance assays

Stationary phase cultures (100 μL) in MP medium (succinate) were used to inoculate 3.5 mL of soft agar (0.8%) that was previously melted and then cooled to ~ 50 °C. Inoculated soft agar was agitated by vortex (~ 10 sec) and then overlaid to solid MP medium (succinate). Soft agar overlays were allowed to solidify at room temperature for 1-2 hr. Antibiotic discs (VWR) saturated with absolute amounts of individual antibiotics (novobiocin (5 μg), tetracycline (30 μg), streptomycin (10 μg), chloramphenicol (30 μg), colistin (10 μg), cefoxitin (30 μg), gentamycin (10 μg), erythromycin (15 μg), rifampicin (5 μg), ciprofloxacin (10 μg), vancomycin (30 μg), kanamycin (30 μg), nalidixic acid (30 μg), ampicillin (10 μg)) were placed on top of agar plates with sterile tweezers. Plates were incubated at 30 °C for ~ 24 hr and then scored for diameter of resulting zones of inhibition.

#### Oxidative stress

Stationary phase cultures were used to prepare soft agar overlays as described for antibiotic resistance assays. After solidification, 5 μL of 30 % hydrogen peroxide (38) was spotted to the center of the plate (38). Plates were incubated at 30 °C for 4-6 d and scored for diameter of resulting zones of inhibition.

#### Alcohol stress

Stationary phase cultures were serially diluted with MP medium (no carbon), plated to solid medium containing 2% ethanol (38) and incubated at 30 °C for 4-6 d and scored for indication of stress (colony size, small or variable) and viability.

#### Heat shock

Mid-exponential phase cultures were placed in a 55 °C water bath for 5 or 10 min (38). Cells were then transferred to room temperature and serially diluted with MP medium (no growth carbon source) and plated to solid medium (succinate). After incubation at 30 °C for 4-6 d they were scored for viability (survival).

### Assessing population heterogeneity

Isolated colonies were inoculated into 2 mL liquid MP media with either 3.5 mM succinate or 15 mM methanol. Upon growth, cultures were diluted 1/64 into 5 mL of the same liquid media in sealed Balch tubes and grown for 24 hr (succinate) or 30 hr (methanol) to allow all cultures to reach stationary phase. After completing this acclimation step, 1 mL of each culture was collected and centrifuged for 1 minute at 10,000xg. The supernatant was removed and the cell pellet was resuspended in 500 μL Diluent C containing 2 μL PKH67 dye (PKH67 Green Fluorescent Cell Linker Kit, Sigma-Aldrich), incubated for 5 minutes at room temperature, and quenched by adding 500 μL 1% BSA solution. Samples were pelleted for 1 minute at 10,000xg and subsequently washed in equal volumes of 1% BSA (once) followed by MP medium (twice). After a final resuspension in an equal volume of MP medium, the final mixes were diluted 1/64 into 5 mL MP media containing 15 mM methanol to initiate experimental cultures and the remainder was mixed with DMSO to 8% and stored at −80 °C. Samples taken during the experimental growth period were also mixed with DMSO to 8% and stored at −80 °C. Signal intensity of samples was assessed via flow cytometry analysis (Beckman Coulter Cytoflex S). After excitation at 488 nm, dye intensity was measured via emission detected with a 525 nm bandpass filter. Cell gating was determined by the distribution of red fluorescence detection from the mCherry-tagged strain CM3839.

## RESULTS

### Loss of a MarR-family regulator confers formaldehyde resistance

To test whether the previously identified *ttmR^EVO^* allele (ref 7) is a loss-of-function allele a precise deletion of the coding region *ttmR* was constructed by allelic exchange in wild-type and its growth was characterized. The one base-pair deletion in *ttmR^EVO^* was in the open reading frame; the frameshift at 140/588 nt yielded a truncated protein with 73 amino acids, only 46 of which were wild-type. The evolved isolate with *ttmR^EVO^* and the *ΔttmR* strain could each use 4 mM formaldehyde as a sole carbon source, reaching a modest absorbance of ~ 0.10 - 0.13 when 4 mM formaldehyde was provided. At 6 mM formaldehyde, both the *ttmR^EVO^* and *ΔttmR* mutants failed to grow after 48 hr, unlike the *ΔefgA* strain (Figures 1A and 2A). Growth in the presence of 4 but not 6 mM formaldehyde was also seen when formaldehyde was added to an additional carbon source, either succinate or methanol (Figure 1BC). In the absence of exogenous formaldehyde (i.e., during growth on succinate or methanol alone), however, the *ΔttmR* strain did not display any differences in lag time, growth rate, or final yield when compared to wild-type (Figure 2B, C, Table S2). Phenotypic comparison of *ttmR* and *ΔefgA* mutants demonstrate that although the *ΔefgA* mutant was more formaldehyde resistant, the mutations are comparable in that they i) allowed formaldehyde growth, ii) did not impact growth in the absence of formaldehyde stress, and ii) conferred formaldehyde resistance during the consumption of alternative carbon sources (Figures 1 and 2) (7).

**Figure 1.**
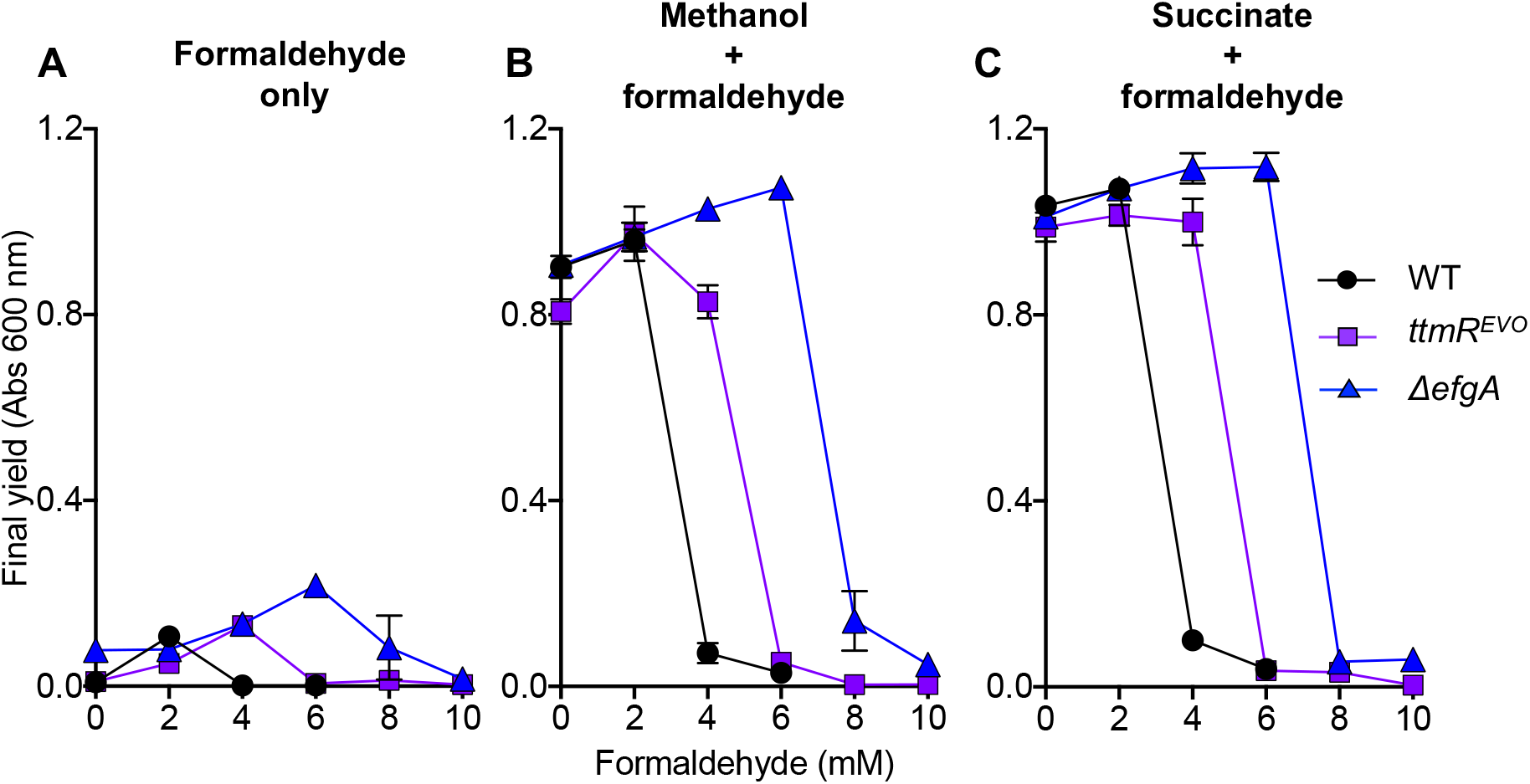
Formaldehyde-evolved isolates have increased formaldehyde resistance. Wild-type (CM2730, black circles) and mutant strains harboring loss-of-function mutations in *ΔefgA* (CM3745, blue triangles) or *ttmR^EVO^* (CM3919, purple squares) were grown in liquid MP medium with 0, 2, 4, 6, 8, or 10 mM exogenous formaldehyde. Final yields are shown when formaldehyde was provided as a sole source of carbon and energy (A) or as a stressor when methanol (B), or succinate (C) was provided as the primary carbon source. *Error bars* represent the standard error of the mean of biological replicates.

**Figure 2.**
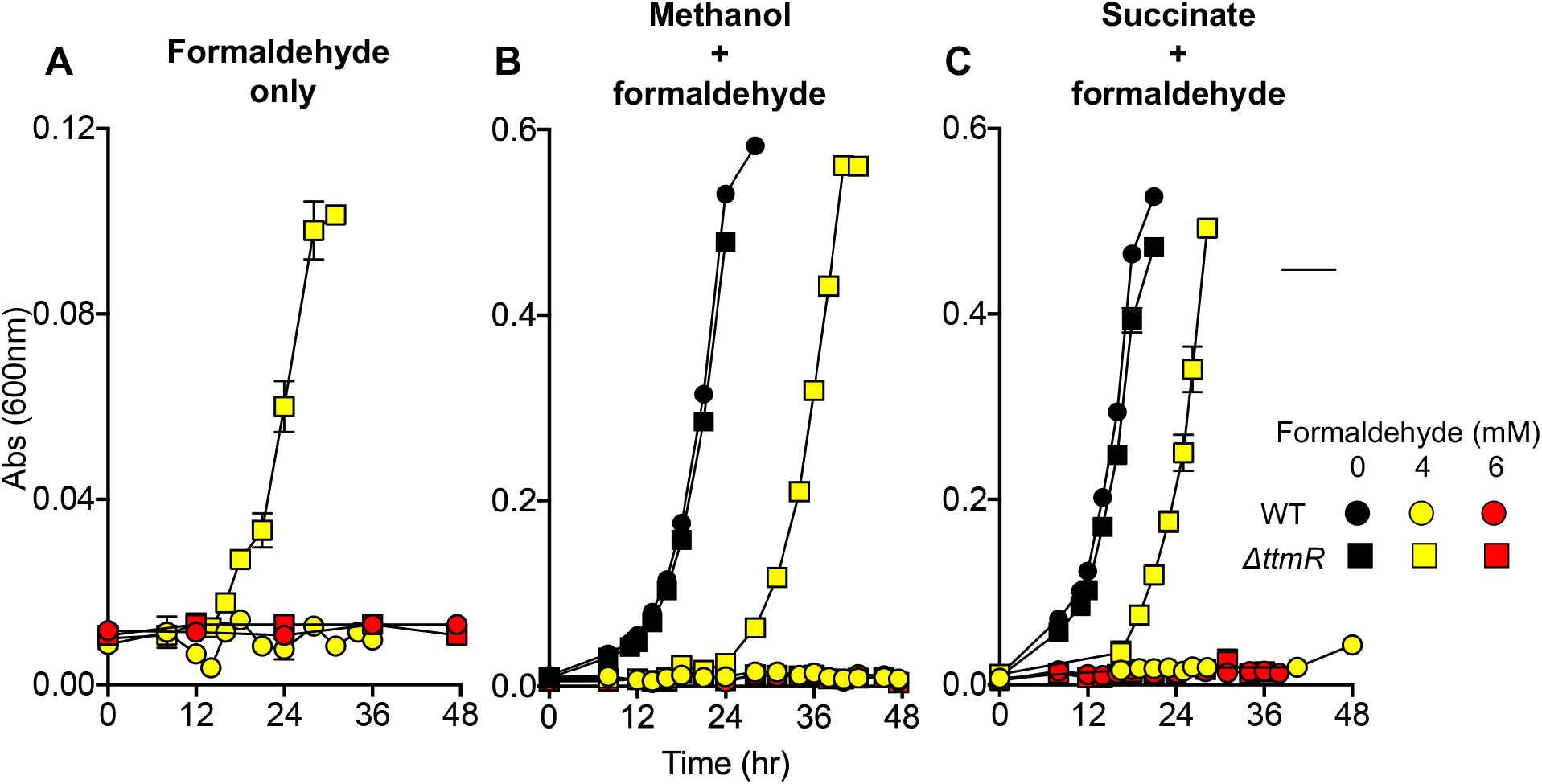
Deletion of *ttmR* recapitulates increased formaldehyde resistance. Wild-type (CM2730, circles) and the constructed *ΔttmR* mutant (CM4732, squares) were grown in liquid MP medium containing 0, 4, or 6 mM exogenous formaldehyde (indicated by black, yellow, or red symbols, respectively). Absorbance up to 48 hours was measured to assess growth when formaldehyde was provided as a sole source of carbon and energy (A) or as a stressor when methanol (B), or succinate (C) was provided as the primary carbon source. *Error bars* represent the standard error of the mean of biological replicates.

### TtmR is not involved in a general stress response

The exact nature of formaldehyde cytotoxicity in *M. extorquens* is unknown; however, in various organisms, previous work has identified numerous forms of formaldehyde-induced DNA damage (39, 40) and more recent work has implicated formaldehyde-induced protein damage in human cell cytotoxicity (41). In considering possible mechanisms of increased formaldehyde resistance, we recognized that cellular processes that are likely to mitigate formaldehyde-induced damage, such as DNA repair and protein quality control, might be involved in generalized stress responses or formaldehyde-specific stress responses. To address whether the loss of TtmR led to a formaldehyde-specific response, the *ttmR^EVO^* strain was exposed to a variety of other stressors. Specifically, we chose stressors that addressed the possibilities that the *ttmR^EVO^* mutant might i) be broadly resistant to aldehydes, ii) confer multidrug resistance (as a homolog of MarR) or iii) mediate a generalized stress response.

Growth assays in media supplemented with various C2 aldehydes (glyoxal and acetaldehyde), as well as a few other aldehydes (butyraldehyde, propionaldehyde, and glutaraldehyde) representing potential metabolic intermediates indicated TtmR appears to be specific to formaldehyde stress. In both succinate and methanol-based media, the *ttmR^EVO^* mutant showed comparable defects in growth to WT other than a very subtle growth improvement when either glyoxal or acetaldehyde was present (Figure S1).

To evaluate antibiotic resistance, disk diffusion assays with different antibiotics were performed with WT or *ΔttmR* mutant soft agar overlays. Wild-type was resistant to 9 of the 14 antibiotics tested (i.e., no zone of inhibition) (Table S3). Of the five remaining antibiotics, the *ΔttmR* mutant did not display any increased resistance (i.e., smaller zone), allowing us to conclude that though there may be differences in specific antibiotic resistances, but there was no broadly acquired antibiotic resistance (Table S3).

Finally, the *ttmR^EVO^* strain was screened for its increased resistance to heat shock (55 °C, treated 24 hr after entry into stationary phase in liquid medium), oxidative stress (H2O2, soft agar overlays), and alcohol stress (EtOH, in solid medium). The *ttmR^EVO^* mutant strain behaved similar to wild-type when exposed to each of the stressors, other than a modest increase in viability during heat shock, which may indicate an elevated protein stress response (Table S3). Collectively, these data demonstrate that TtmR is not a general stress response protein but rather its activity is specific to modulating stress induced by formaldehyde.

### TtmR modulates the expression of genes involved in an array of cellular functions

To understand the impact of TtmR on the transcriptome, we performed RNA-sequencing analysis on wild-type and the *ΔttmR* mutant during early exponential growth (OD = 0.2) in minimal medium with 15 mM methanol. Specifically, we aimed to identify differentially expressed genes (DEGs) that would explain the increased resistance to formaldehyde.

A pair-wise comparison of log-transformed counts of wild-type and the *ΔttmR* mutant was performed and DEGs were identified by imposing a two-fold change cutoff (Log_2_FC > 1.0) with an adjusted p-value (p_adj_) less than 0.05. In total, we identified 75 DEGs in the *ΔttmR* mutant, of which 61 (81%) were upregulated and 14 (19%) were downregulated (Tables 1, Figures 3 and S2). The expression differences observed in the upregulated genes were quite large; 34 had fold changes ≥4X (Log_2_FC>2.0) and six had fold changes ≥10X (Log_2_FC≥3.3). By contrast, all downregulated genes had ≤4.6X fold change. Notably, several of the most upregulated DEGs (eg, 5 of the 6 with fold changes ≥10X) were clustered in the chromosome in operons or multiple adjacent operons (Figure 3).

**Figure 3.**
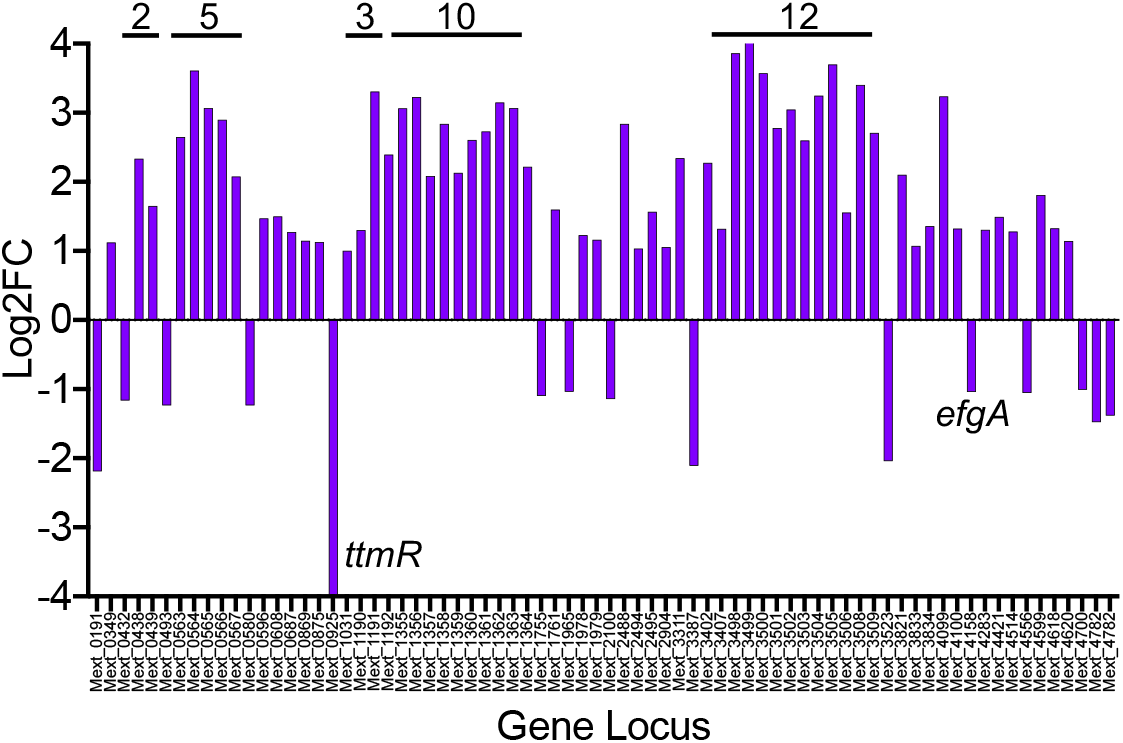
Differentially expressed genes in the *ΔttmR* mutant. RNA-sequencing analysis was performed on wild-type and the *ΔttmR* mutant with cells in the early exponential phase of growth on methanol. The log_2_ fold changes (Log_2_FC) in gene expression of the *ΔttmR* mutant were calculated relative to wild-type. Adjusted p-values were also calculated. Differentially expressed genes are defined as having a Log_2_FC > 1.0 and p_adj_ < 0.05; each bar represents an individual gene. Horizontal bars over clusters of genes indicate two or more adjacent genes with the number of genes clustered indicated above the bar. Gene annotations are provided in Table 1.

**Table 1.** Genes differentially expressed in a *ΔttmR* mutant. RNA-sequencing analysis was performed on wild-type and the *Δ ttmR* mutant with cells in the early exponential phase of growth on methanol. The log_2_ fold changes (Log_2_FC) in gene expression of the Δ *ttmR* mutant were calculated relative to wild-type. Adjusted p-values were also calculated. Genes detailed here have a Log_2_FC > 1.0 and p_adj_ < 0.05.

| Locus | Annotation (RefSeq) | Functional category | Log<sub>2</sub>FC | p<sub>adj</sub> |
| --- | --- | --- | --- | --- |
| Mext_0044 | hypothetical protein | Hypothetical | 1.015 | 3.00E-3 |
| Mext_0191 | chemotaxis sensory transducer | Chemotaxis | -2.189 | 2.81E-129 |
| Mext_0349 | flagellin domain protein | Motility | 1.121 | 8.86E-17 |
| Mext_0432 | ROSMUCR transcriptional regulator | Regulatory | -1.166 | 8.85E-44 |
| Mext_0438 | hypothetical protein | Signaling | 2.335 | 1.36E-143 |
| Mext_0439 | hypothetical protein | Regulatory | 1.652 | 1.24E-25 |
| Mext_0493 | hypothetical protein | Inclusion bodies | -1.235 | 2.44E-16 |
| Mext_0563 | acetate kinase | Central carbon metabolism | 2.647 | 1.15E-124 |
| Mext_0564 | secretion protein HlyD family protein | Transport | 3.610 | 0.00 |
| Mext_0565 | ABC transporter related | Transport | 3.068 | 6.09E-291 |
| Mext_0566 | ABC-2 type transporter | Transport | 2.898 | 1.08E-144 |
| Mext_0567 | Phosphoketolase | Central carbon metabolism | 2.076 | 4.73E-148 |
| Mext_0580 | pseudogene | Pseudogene | -1.234 | 8.45E-38 |
| Mext_0596 | response regulator receiver | Regulatory | 1.472 | 3.29E-81 |
| Mext_0608 | helix-turn-helix- domain containing protein AraC type | Regulatory | 1.501 | 1.27E-44 |
| Mext_0687 | flagellar FlbT family protein | Motility | 1.274 | 3.95E-35 |
| Mext_0869 | hypothetical protein | Stress (oxidative) | 1.149 | 5.60E-10 |
| Mext_0875 | PAS sensor protein | Signaling (heat shock) | 1.131 | 5.67E-37 |
| Mext_0925 | regulatory protein MarR | MarR | -9.624 | 1.38E-95 |
| Mext_1031 | glycerone kinase | Central carbon metabolism | 1.002 | 6.93E-21 |
| Mext_1190 | PAS sensor protein | Signaling | 1.304 | 1.92E-64 |
| Mext_1191 | membrane protein of unknown function UCP014873 | Membrane DUF | 3.307 | 0.00 |
| Mext_1192 | 67 kDa myosin-cross-reactive antigen family protein | Stress (FA metabolism) | 2.395 | 1.81E-143 |
| Mext_1355 | UspA domain protein | Stress | 3.064 | 0.00 |
| Mext_1356 | cytochrome c class I | Cytochrome metabolism | 3.227 | 0.00 |
| Mext_1357 | cyclic nucleotide-binding | Regulatory (cNMP) | 2.086 | 6.65E-176 |
| Mext_1358 | ABC transporter related | Transport | 2.837 | 2.01E-249 |
| Mext_1359 | ABC transporter related | Transport | 2.131 | 9.30E-160 |
| Mext_1360 | cytochrome bd ubiquinol oxidase subunit I | Cytochrome metabolism | 2.607 | 1.66E-216 |
| Mext_1361 | cytochrome d ubiquinol oxidase, subunit II | Cytochrome metabolism | 2.728 | 2.92E-238 |
| Mext_1362 | *cyd* operon protein YbgT | Cytochrome metabolism | 3.150 | 2.69E-256 |
| Mext_1363 | regulatory protein DeoR | Regulatory (sugar) | 3.069 | 0.00 |
| Mext_1364 | amidase | Translation (tRNA charging) | 2.218 | 7.97E-68 |
| Mext_1755 | aminoglycoside phosphotransferase | Signaling (phosphotransferase) | -1.098 | 7.26E-27 |
| Mext_1761 | methionine-*R*-sulfoxide reductase | Stress (oxidative), Regulatory (PTM) | 1.598 | 2.08E-37 |
| Mext_1965 | response regulator receiver | Signaling | -1.037 | 8.82E-35 |
| Mext_1978 | diguanylate cyclase | Signaling (c-di-GMP) | 1.230 | 1.73E-37 |
| Mext_1979 | hypothetical protein | hypothetical | 1.160 | 1.62E-08 |
| Mext_2100 | hypothetical protein | Hypothetical (transport) | -1.145 | 4.33E-38 |
| Mext_2114 | hypothetical protein | Hypothetical | 1.186 | 6.88E-3 |
| Mext_2488 | OsmC family protein | Stress (oxidative) | 2.837 | 2.28E-184 |
| Mext_2494 | conserved hypothetical membrane spanning protein | Hypothetical (membrane) | 1.035 | 2.75E-10 |
| Mext_2495 | CsbD family protein | Stress | 1.568 | 2.67E-19 |
| Mext_2904 | hypothetical protein | Regulatory (stress) | 1.055 | 2.33E-10 |
| Mext_3311 | hypothetical protein | Hypothetical | 2.342 | 4.15E-117 |
| Mext_3387 | hypothetical protein | Hypothetical | -2.109 | 2.83E-46 |
| Mext_3402 | hypothetical protein | Hypothetical | 2.275 | 2.38E-121 |
| Mext_3407 | ABC transporter related | Transport | 1.319 | 3.12E-06 |
| Mext_3498 | heat shock protein Hsp20 | Chaperone/Heat shock | 3.861 | 0.00 |
| Mext_3499 | putative transcriptional regulatory protein, Crp/Fnr family | Regulatory (cNMP) | 4.117 | 0.00 |
| Mext_3500 | UspA domain protein | Stress (universal) | 3.573 | 0.00 |
| Mext_3501 | hypothetical protein | Translation (ribosomal conflict) | 2.779 | 1.99E-260 |
| Mext_3502 | transport-associated | Transport | 3.045 | 0.00 |
| Mext_3503 | hypothetical protein | Hypothetical | 2.599 | 1.25E-148 |
| Mext_3504 | UspA domain protein | Stress (universal) | 3.249 | 0.00 |
| Mext_3505 | regulatory protein Crp | Regulatory (cNMP) | 3.699 | 0.00 |
| Mext_3506 | response regulator receiver | Regulatory | 1.557 | 9.75E-27 |
| Mext_3508 | UspA domain protein | Stress (universal) | 3.401 | 0.00 |
| Mext_3509 | metal-dependent phosphohydrolase HD sub domain | Signaling | 2.707 | 2.11E-242 |
| Mext_3523 | hemolysin-type calcium-binding region | C-N hydrolase | -2.042 | 8.32E-29 |
| Mext_3821 | choloylglycine hydrolase | C-N hydrolase | 2.101 | 1.03E-99 |
| Mext_3833 | cytochrome d ubiquinol oxidase, subunit II | Cytochrome metabolism | 1.073 | 7.21E-17 |
| Mext_3834 | cytochrome bd ubiquinol oxidase subunit I | Cytochrome metabolism | 1.358 | 2.35E-32 |
| Mext_4099 | ATP-binding region ATPase domain protein | Signaling | 3.238 | 0.00 |
| Mext_4100 | carbonic anhydrase | pH | 1.323 | 4.67E-28 |
| Mext_4158 | EfgA | Stress | -1.041 | 5.94E-35 |
| Mext_4283 | hypothetical protein | Hypothetical | 1.306 | 6.29E-15 |
| Mext_4421 | hypothetical protein | Hypothetical | 1.492 | 4.11E-24 |
| Mext_4514 | autoinducer-binding domain protein | Regulatory (quorum sensing) | 1.280 | 2.40E-39 |
| Mext_4556 | heat shock protein Hsp20 | Chaperone/heat shock | -1.055 | 8.33E-33 |
| Mext_4599 | flavin reductase domain protein FMN-binding | Redox rxn | 1.809 | 8.86E-10 |
| Mext_4618 | nucleotide sugar dehydrogenase | Sugar metabolism | 1.325 | 7.69E-05 |
| Mext_4620 | NAD-dependent epimerase/dehydratase | Sugar metabolism | 1.145 | 6.81E-13 |
| Mext_4700 | integrase, catalytic region | Phage | -1.009 | 1.14E-05 |
| Mext_4782 | chaperonin GroEL | Chaperone/Heat shock | -1.477 | 6.20E-74 |
| Mext_4783 | chaperonin Cpn10 | Chaperone/Heat shock | -1.385 | 4.09E-57 |

Using their RefSeq annotations and Pfam domains, gene functionality of all of the DEGs were categorized into 14 groups which included 12 groups of genes with inferred functionality (n=58), one group that encoded hypothetical proteins (n=11), and one group of singletons (n=6), whose functionality was only represented by a single DEG (Figure S3). Of the 58 DEGs with a proposed function, the largest categories were Regulatory (n=12), Stress (n=10), Signaling (n=8), and Transport (n=7). Genes classified as Regulatory were comprised of a variety of transcription factors, sigma factors, or anti-sigma factors. Of the Regulatory DEGs, three were Crp homologs, predicted to bind cyclic nucleotides, and three were predicted to have roles in stress response (oxidative stress, cold shock, and generalized stress response, respectively). As many of the DEGs are themselves regulatory, the full array of DEGs identified in the *ΔttmR* mutant likely represent loci that are directly controlled by TtmR and indirectly controlled by TtmR, through other regulators. The Signaling group included genes that encoded one or more PAS sensor domains and/or predicted members of two-component regulatory systems such as histidine kinases or response regulator receivers (single domain or paired with diguanylate cyclase domain). DEGs of the Stress group encoded universal stress proteins, whose mechanisms are highly variable, and proteins predicted to be involved in oxidative stress and solvent stress. Genes in the Transport grouping were involved in a Type I Secretion System (T1SS) and ABC transporters. There was little indication about the molecules that might be transported, with the exception of an annotated sulfonate transporter. The remaining DEGs were involved in Carbon metabolism, Cytochrome metabolism, C-N hydrolase and Chaperone/Heat shock.

The DEGs that were downregulated in the *ΔttmR* mutant strain were scattered across functional categories with the exception of three of the four DEGs encoding Chaperonin/Heat shock proteins. One possibility for the downregulated DEGs is that they are positively regulated by TtmR, when it is present.

Interestingly, we noticed inconsistency in the directionality of expression changes for Chaperone genes. Chaperonin GroEL, Chaperonin Cpn10, and Heat shock protein Hsp20 were downregulated while a second Heat shock protein Hsp20 homolog, *Mext_3498,* was significantly upregulated (14.5X, Log_2_FC=3.9). *Mext_3498* was part of the highly upregulated twelve-gene cluster and appears to be the first gene in a two-gene operon that also encoded a Crp homolog with a cyclic nucleotide binding motif. These non-uniform changes may reflect the distinct roles of Chaperones involved in housekeeping protein quality control versus those which are stress-related. The considerable upregulation of Heat shock protein Hsp20 homolog, *Mext_3498*, may prevent formaldehyde-induced protein damage even in the midst of downregulation of Chaperonin GroEL, Chaperonin Cpn10, and a second homolog of Heat shock protein Hsp20.

Of the 75 DEGs identified herein, 20 of them were identified in the 33 previously associated with the transcriptome of the phenotypically formaldehyde-tolerant subpopulation of wild-type *M. extorquens* described elsewhere (42). We find that the directionalities, but not the magnitudes, of expression changes are comparable to those found in the formaldehyde-tolerant subpopulation (Figure 3, Table 1). In all 20 instances, gene expression differences were more pronounced in the *ΔttmR* mutant than in the subpopulation. The genes of the methanol utilization pathways in the *ΔttmR* mutant did not meet our conservative criteria for being identified as DEGs. However, closer examination showed modest decreases (~ 1.3-1.6 fold changes, p-adj < 0.05) in the expression of *mxaF,* which encodes the large subunit of methanol dehydrogenase (43, 44), as well as genes involved in formate assimilation and oxidation (Table S4).

Overall our data suggests that TtmR elimination allows formaldehyde resistance by tuning the expression of a number of loci that collectively launch a formaldehyde-specific physiological response that may include downstream regulation, cyclic nucleotide signaling, transport, and general stress responses. Notably, the DEGs identified were not involved in cellular functions most often associated with formaldehyde stress, such as DNA repair or formaldehyde detoxification by methylotrophic pathways or alcohol/aldehyde dehydrogenases.

### TtmR can modulate formaldehyde resistance independent of EfgA

EfgA is a predicted formaldehyde sensor that modulates formaldehyde resistance by halting translation in the presence of elevated formaldehyde (7). Like TtmR, the absence of EfgA confers *M. extorquens* with the ability to grow on formaldehyde as a sole source of carbon and energy (Figure 1A), presumably because formaldehyde-mediated translational pausing has been eliminated. Our transcriptomic analysis showed that *efgA* was ~ 2.0X downregulated in the *ΔttmR* mutant. Thus, our data suggested it was possible that TtmR mediates formaldehyde resistance through regulating *efgA* expression.

To test this hypothesis, we constructed a *ΔttmR ΔefgA* double mutant by successive allelic exchange. We then compared the growth of the double mutant to that of each of the single mutants with variable concentrations of formaldehyde provided in the media. In the absence of exogenous formaldehyde, growth of the double mutant was indistinguishable from either of the single mutants (data not shown). By contrast, when 6 mM formaldehyde was added to the growth medium, the *ΔttmR* mutant failed to grow after 24 hr, while the *ΔttmR ΔefgA* double mutant was more fit than the *ΔefgA* single mutant (Figure 4). These data demonstrated that TtmR has some effects that extend beyond regulating EfgA, although the transcriptomic data suggest that the formaldehyde resistance of *ttmR* mutants might be partially mediated by decreased *efgA* expression.

**Figure 4.**
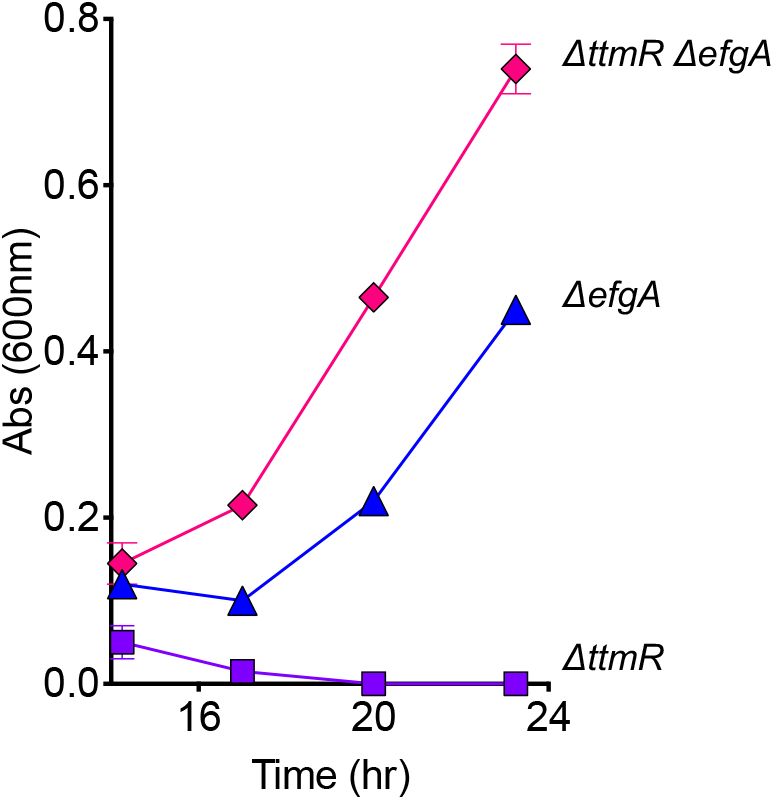
The *ΔttmR ΔefgA* double mutant has increased formaldehyde resistance. Growth of the *ΔttmR* (CM4732, purple squares), *ΔefgA* mutant (CM3745, blue triangles), and *ΔttmR ΔefgA* double mutant (CM4733, pink diamonds) was quantified in liquid MP medium (succinate) containing 6 mM formaldehyde. *Error bars* represent the standard error of mean of biological replicates.

### TtmR and EfgA are required for optimal transition to methylotrophy

To understand when TtmR and EfgA would be beneficial to *M. extorquens*, we sought to identify growth conditions that induced formaldehyde stress in a physiologically relevant context (i.e., from methanol utilization). Previous work has demonstrated that during the switch from multi-carbon to single-carbon growth, *M. extorquens* AM1 experiences a transient formaldehyde imbalance, such that formaldehyde is secreted into the growth medium (45). Researchers suggested that imbalance occurs as a result of non-transcriptional regulation preventing flux through methylotrophic pathways where toxic metabolites (formaldehyde, glycine and glyoxylate) are produced.

We assayed growth of wild-type, *ΔttmR,* and *ΔefgA* strains during the transition to methylotrophy by subculturing stationary-phase, succinate-grown cells into fresh medium containing methanol as the sole carbon source. Both mutant strains displayed lag times that were comparable to each other but significantly longer (by ~ 6 hr) than those observed in wild-type (Table 2, Figure 5A). However, the growth rates and final yields of all strains were indistinguishable (Table 2). Cell viability assays uniformly showed that there was no statistically significant decrease in viable cell counts (WT vs. *ΔttmR* p-value=0.223, WT vs. *ΔefgA* p-value=0.215) during the apparent lag phases and indicated that mutant strains take longer to enter exponential growth, rather than experiencing death (Figure 5C). The ability of strains to transition between distinct modes of metabolism was further probed in an analogous experiment, where methanol-grown cells (Figure 5A) were subcultured into fresh medium containing succinate as the sole carbon source; no growth defect was observed (Figure 5B).

**Figure 5.**
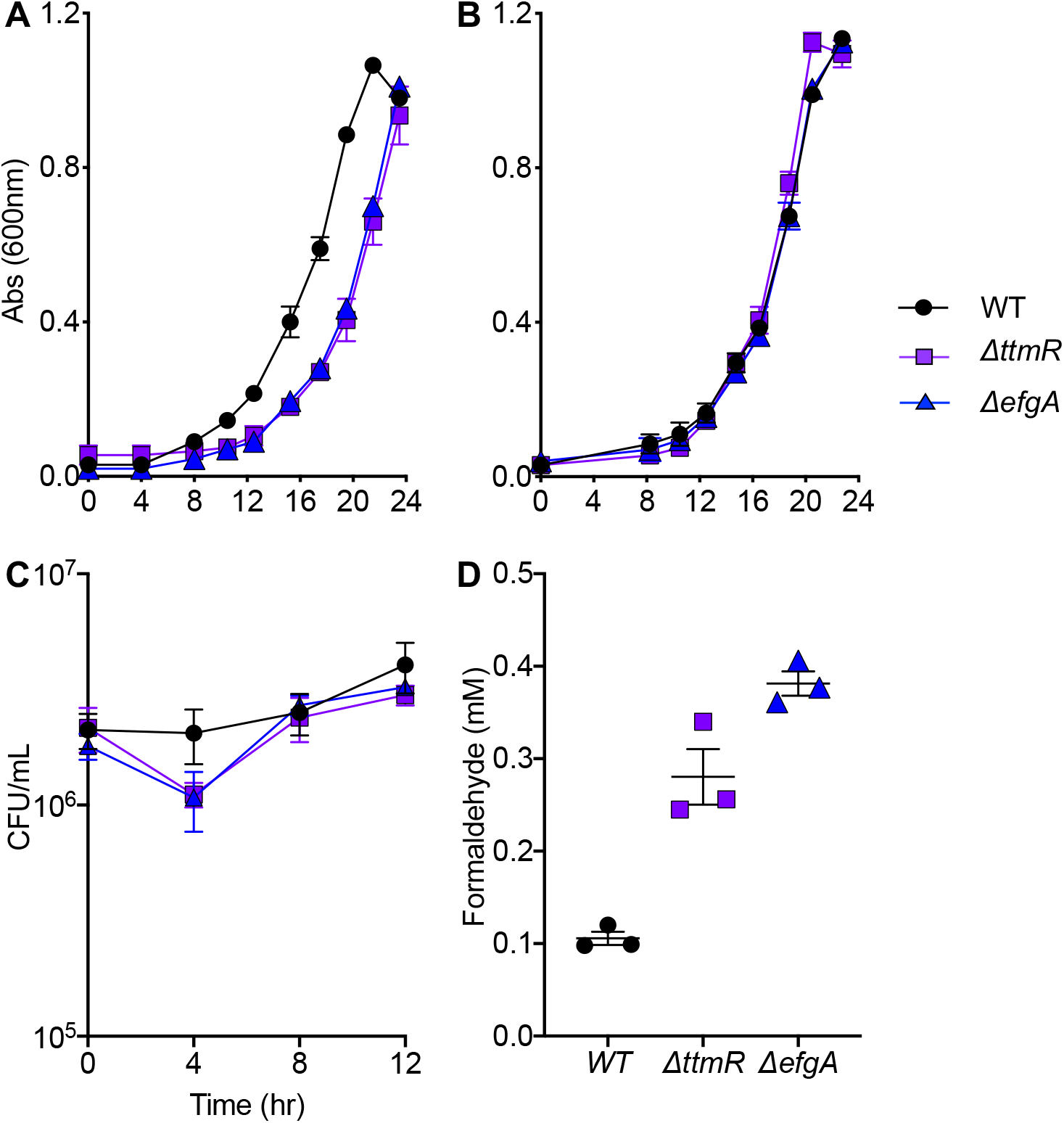
The *ΔttmR* and *ΔefgA* mutants are defective in the transition to methylotrophy. The wild-type (CM2730, black circles), *ΔttmR* mutant (CM4732, purple squares), and *ΔefgA* mutant (CM3745, blue triangles) were first acclimated to succinate in liquid MP medium. Their growth was then assayed upon their inoculation to methanol-based medium by absorbance (A) and cell viability (C). Formaldehyde concentrations in the growth medium was measured by a colorimetric assay (D). After reaching stationary phase, the methanol-grown cultures were subcultured to succinate-based medium and assayed for growth by absorbance (B). We used a >15% increase from starting absorbance (dashed line) constituted a threshold that marked the end of lag phase. *Error bars* represent the standard error of mean of biological replicates. In panel D, individual values are shown and the mean is indicated.

**Table 2.** Growth during carbon substrate transitions. Growth of succinate- or methanol-grown stationary phase cells subcultured into fresh medium with 15 mM methanol or 3.5 mM succinate, respectively, was monitored. Error shown represents standard error of the mean of biological replicates. Error was not provided for lag time as resolution at low cell densities is not sensitive enough to quantify. A two-tailed Student’s t-test was performed to identify statistically significant differences in final yields or growth rates between growth rates (p-value < 0.05).

| Switch | Strain |  | Lag (hr) | Final Yield (Abs600) | p-value | Growth rate<br>(μ) | p-value |
| --- | --- | --- | --- | --- | --- | --- | --- |
| S → M | WT |  | 8 | 0.98 +/- 0.028 |  | 0.2139 +/- 0.002 |  |
|  | *ΔttmR* |  | 14 | 0.84 +/- 0.014 | 0.090 | 0.2295 +/- 0.011 | 0.031 |
|  | *ΔefgA* |  | 14 | 0.96 +/- 0.000 | 0.493 | 0.2077 +/- 0.001 | 0.575 |
| M → S | WT |  | 8 | 1.13 +/- 0.000 |  | 0.2333 +/- 0.018 |  |
|  | *ΔttmR* |  | 8 | 1.13 +/- 0.000 | 0.500 | 0.2491 +/- 0.003 | 0.471 |
|  | *ΔefgA* |  | 8 | 1.12 +/- 0.014 | NV | 0.2325 +/- 0.018 | 0.960 |

To further characterize the defect in the transition to methylotrophy of the *ΔttmR* and *ΔefgA* strains, we monitored formaldehyde accumulation in the culture supernatants. In wild-type, formaldehyde accumulated in the supernatant and peaked at ~ 100 μM upon entry into exponential growth, comparable to previously described results in *M. extorquens* AM1 (35). In the supernatants of the *ΔttmR* and *ΔefgA* mutants, formaldehyde accumulated to nearly 3-4X the levels observed in wild-type (~ 275 and ~ 400 μM, respectively) demonstrating that formaldehyde resistant mutants have perturbed formaldehyde metabolism (Figure 5D).

Collectively, these data show that during the transition to methylotrophy, formaldehyde imbalance is exacerbated in the absence of TtmR or EfgA and suggest that both proteins are required for formaldehyde homeostasis. Therefore, in mutant strains, there is a clear tradeoff between acquiring formaldehyde resistance and the ability to adapt to methylotrophic growth.

### Phenotypic heterogeneity during the transition to methylotrophy

It has been previously observed that bacterial stressors can provoke phenotypic heterogeneity and, recently, phenotypic heterogeneity in formaldehyde tolerance was demonstrated in wild-type populations of unstressed *M. extorquens* PA1 (42). Here, in the presence of elevated formaldehyde, individual cells experience a binary outcome and either tolerate formaldehyde from the onset of exposure and grow or succumb to formaldehyde stress and die. As the transition to methylotrophy leads to formaldehyde imbalance, we wanted to determine if the extended lag times of the *ΔttmR* and *ΔefgA* mutants during this transition were due to increased variance of individuals entering exponential growth (i.e., some cells initiating growth after a longer time or not at all) or was indicative of a change in the mean behavior of individuals without increased variance.

Tracking the entry into growth in populations by dilution of a membrane-intercalating dye in individual cells revealed heterogeneous initiation of growth in *ΔttmR* and *ΔefgA* strains. When cells were transferred from one methanol-grown culture to fresh medium with methanol, all strains displayed a gradual and uniform lessening in the distributions of per-cell fluorescence, indicating uniform growth of cells within these populations (Figures 6 and S4). In contrast, following the transition to methylotrophic growth, a notable skew, or split, in the distribution was observable by 9 hours. Most pronounced in the *ΔefgA* mutant, this result suggests the existence of a subpopulation of nongrowing cells in the *ΔefgA* lineages, which is not effectively transitioning to growth on methanol. The *ΔttmR* lineages experience a similar delay; however, in this genotype, the effect is seen as a rightward skew of the distribution rather than as two distinct peaks, suggesting that not all cells survive the transition. This effect was dramatically diminished in the wild-type strain, suggesting that it experiences more uniformity and less growth inhibition during the transition between growth on different carbon substrates across members of the population. This result is also consistent with bulk growth trends observed in the methylotrophic switch (Figure 5).

**Figure 6.**
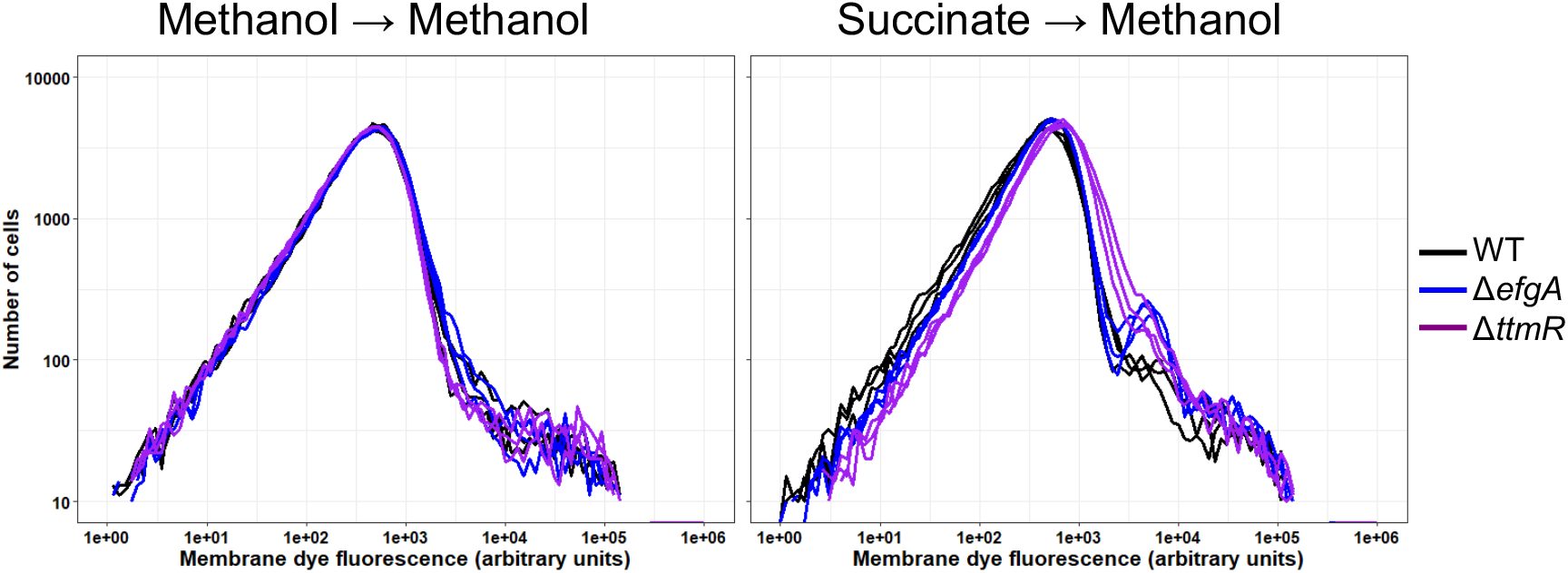
Distribution of cell division during growth on methanol varies depending on the starting growth substrate. Following acclimation to growth on either single- (methanol) or multi-carbon (succinate) sources, stationary phase cells were labelled with a fluorescent dye that intercalates into the cell membrane and thereby allows tracking of cell division through dilution of the starting signal. Patterns of cell division were measured following 24 hours of growth in methanol (A) or during shifts from succinate to methanol (B) by measuring fluorescence of individual cells within the populations using flow cytometry. The resulting data is displayed as distributions of fluorescence intensity that demonstrate how a population does or does not migrate (grow and divide) over time. Data is plotted as histograms of per-cell fluorescence intensity (measured at the emission wavelength of 525 nm, FITC channel) of cell populations at different time points across growth. Each line in a panel represents cell events surveyed for a single (of three) biological replicate within an equivalent number of cells.

## DISCUSSION

During methylotrophic growth, all carbon flows through formaldehyde, a cytotoxic compound that can potentially damage myriad biomolecules. Accordingly, methylotrophs are more resistant to formaldehyde than non-methylotrophs (42). In large part, formaldehyde resistance can be attributed to the constitutively high expression and activity of Fae, which catalyzes the first step of formaldehyde oxidation and eliminates free formaldehyde from the cytoplasm. Despite this, formaldehyde pools in *M. extorquens* are elevated in comparison to non-methylotrophs (1, 46–49) and the mechanisms by which these greater pool sizes are tolerated and regulated are largely unknown, as are specific mechanisms of averting formaldehyde-induced cellular damage. Recent work uncovered the role of EfgA in directly sensing formaldehyde and acting to inhibit translation (7). The current work indicates that TtmR, a MarR-like transcription factor, is a second regulator in the formaldehyde stress response predicted to act at the level of transcription. TtmR regulates transcription of *efgA*, but the growth phenotypes of double mutants also indicate that TtmR regulates other targets beyond this. TtmR and EfgA each contribute to formaldehyde homeostasis and are particularly critical during the transition to methylotrophic metabolism. The combination of these distinct systems suggests that formaldehyde metabolism is monitored and managed across transcriptional and translational regulatory processes.

Without TtmR, *M. extorquens* does not display any obvious growth defects during methylotrophic growth, but instead, displays increased formaldehyde resistance. Increased formaldehyde resistance, as measured by growth in the presence of increasing formaldehyde concentrations, can be achieved by alleviating formaldehyde damage or by relaxing regulatory mechanisms that would otherwise prevent growth in potentially harmful conditions. Characterized formaldehyde stress response systems in bacteria prevent formaldehyde damage by formaldehyde detoxification and are analogous to formaldehyde oxidation systems used by a variety of methylotrophs for C_1_ utilization (21). In many organisms where formaldehyde stress has been investigated, formaldehyde-induced DNA damage is often identified and cell damage and death are attributed to genotoxicity (39, 40, 50). More recently however, formaldehyde-induced proteotoxicity has been implicated in cytotoxicity in some human tissues (41), in synthetic methylotrophs (51), and has been suggested by our own work in *M. extorquens* (7, 42). Therefore, other mechanisms of formaldehyde resistance might activate appropriate DNA repair systems and/or heat shock proteins to mitigate DNA and/or protein damage.

Surprisingly, in the transcriptome of the *ΔttmR* mutant there are no DEGs that are obviously linked to methylotrophic pathways, independent formaldehyde detoxifying enzymatic activities, or DNA repair. Further, of the four genes grouped into Chaperone/Heat shock, three of them are downregulated in the absence of TtmR. Functional groupings showed that the majority of DEGs encode regulators, signaling proteins, stress response proteins, hypothetical proteins, and transporters whose functions and physiological roles are unknown. This variety of DEGs suggests that formaldehyde resistance may be the result of cumulative, coordinated responses from many uncharacterized mechanisms.

Although increased formaldehyde resistance that emerges in strains lacking TtmR or EfgA could be an advantageous trait in a methylotroph, we have demonstrated that a physiological and ecological benefit of TtmR and EfgA is to speed the transition to methylotrophy when endogenous formaldehyde is known to accumulate. Many *Methylobacterium/Methylorubrum* are plant-associated and can use methanol derived from plants by pectin metabolism, among other compounds (26, 52–56). Some, such as *M. extorquens* PA1, which was isolated from *Arabidopsis thaliana*, are epiphytes and live on plant leaf surfaces (26). On leaf surfaces, methanol is released from stomata, small pores in plant tissues that allow gas exchange (57). Methanol availability, therefore, fluctuates as stomata open and close and is most abundant in the morning, upon the first stomatal opening of the day (58, 59). To take advantage of this nutritional niche, *M. extorquens* must quickly transition to methylotrophic metabolism, which necessarily includes production of a metabolic stressor, formaldehyde (1, 45). Therefore, during this transition the physiological state of *M. extorquens* must also be prepared for formaldehyde stress to occur. Here, TtmR and EfgA were shown to be required for the optimal transition to methylotrophic growth and further, in their absence, excess formaldehyde accumulation was observed, demonstrating dysregulation of formaldehyde metabolism. On the phylloplane, the organism’s native environment, where methanol emissions are transient and come in bursts, loss of TtmR or EfgA would likely be disadvantageous and render *M. extorquens* less competitive within its ecological niche where methanol capture is important.

The transition to methylotrophy in *M. extorquens* AM1 has been previously investigated with systems biology and genetic approaches, but did not identify either the *ttmR* or *efgA* locus (1, 5, 6, 45). There, investigators found that upon transitioning from the multi-carbon growth substrate succinate to the single-carbon growth substrate methanol, cells have a lag in carbon assimilation and experience a transient imbalance in formaldehyde metabolism, with formaldehyde accumulating such that it was secreted into the growth medium (45). Further investigation revealed that in adapting cells, carbon flux restriction was mediated by methenyl-dH4MPT, an intermediate in the formaldehyde dissimilation that directly inhibits MtdA, the enzyme that generates the substrate for entry into the Serine Cycle assimilation pathway and may regulate Fae activity by an unknown mechanism (1). These authors propose carbon flux restriction is an adaptive strategy that minimizes the accumulation of toxic metabolites generated in methylotrophic pathways, including formaldehyde. As strains defective in dH4MPT biosynthesis *(ΔmptG)* were previously shown to be more methanol (i.e., formaldehyde) sensitive in the absence of EfgA (7), we conclude that EfgA-mediated formaldehyde homeostasis does not involve methenyl-dH4MPT.

All of the phenotypes of the *ΔttmR* mutant were mirrored in the *ΔefgA* mutant; therefore, we considered the possibility that TtmR and EfgA were interconnected. Specifically, we considered whether TtmR was a positive regulator of *efgA* expression. The transcriptomic data initially seemed to support this possibility, as *efgA* expression was two-fold lower in the *ΔttmR* mutant. However, genetic analysis showed that TtmR and EfgA elimination could cause formaldehyde resistance and a defective switch to methylotrophy independent of one another. Thus, this work has identified TtmR as a second formaldehyde stress response protein, which can act independently of EfgA and a role for TtmR and EfgA in formaldehyde tolerance and homeostasis. Given the phenotypic overlap of these proteins, it is still possible, and perhaps quite likely, that they independently impact a common cellular function that results in the formaldehyde-associated phenotypes observed. Alternatively, the overlap may represent that *efgA* expression is just one of multiple mechanisms that TtmR exerts its effects on formaldehyde homeostasis.

Herein, we demonstrate that achieving formaldehyde resistance through the loss of TtmR or EfgA comes at a cost, as increased formaldehyde resistance simultaneously limits the adaptability of *M. extorquens* to methylotrophic metabolism. The formaldehyde accumulation observed during the transition to methylotrophy suggest that strains lacking TtmR and EfgA achieve formaldehyde resistance by losing their ability to sense and respond to perturbations in formaldehyde homeostasis, rather than by simply improving cellular processes that reduce formaldehyde damage. Interestingly, this tradeoff can be avoided, as mutations that permitted growth on formaldehyde that occurred in other loci, *def* and *efgB*, did not cause a defect in the transition to methylotrophy (Figure S5). Therefore, the transcriptomic profile observed in strains lacking TtmR likely reflects relaxing a combination of cellular components that typically govern formaldehyde balance as well as those that are responding to the resulting formaldehyde perturbation. The variety of DEGs, which includes many regulators, suggest that TtmR is part of a larger gene regulatory network. Additionally, it appears that both TtmR and EfgA are required for *M. extorquens* to rapidly shift from growth on multi-carbon substrates to growth on one-carbon substrates and may be of general importance for facultative methylotrophs.

**Figure S1. Treatment of a *ΔttmR* mutant with alternative aldehydes.**

Growth of wild-type (CM2730, black circles) and the *ttmR^EVO^* mutant (CM3919, purple squares) was quantified in liquid MP medium (succinate) with the addition of (A) 2 mM formaldehyde, (B) 1.25 mM acetaldehyde, (C) 2.5 mM butyraldehyde, (D) 2.5 mM propionaldehyde, (E) 1.25 mM glyoxal, and (F) 0.157 mM glutaraldehyde. Growth of wild-type was quantified in the same medium (succinate) without aldehydes and is superimposed in all panels for reference (CM2730, gray circles). Growth of CM3919 was indistinguishable from wild-type in the absence of aldehyde stress; data were excluded for visual simplicity. *Error bars* represent the standard error of mean of biological replicates.

**Figure S2. Histogram of adjusted p-values and Log_2_FC values for RNA-Seq.**

The distributions of calculated (A) p-values and (B) Log_2_FC values are shown. Bins are defined in increments of 0.05. The red line shows the curve fit to the Gaussian distribution of Log_2_FC values.

**Figure S3. Distribution of DEGs in different cellular processes.**

The pie graph shows the distribution of differentially expressed genes in functional categories according to cellular process. Categories were assigned by annotation and inference from the presence of specific Pfam domains. When function could not be inferred (lack of Pfam domain, domain of unknown function), genes were classified as Hypothetical (orange) and when function was only represented once, genes were classified as Singletons (white). Number of genes in each category is indicated as well as the proportion are up- or down-regulated (arrows).

**Figure S4. Heterogeneity during the transition to methylotrophic metabolism.**

Following acclimation to growth on either single- or multi-carbon sources, stationary phase cells were labelled with a fluorescent dye that intercalates into the cell membrane and thereby allows tracking of cell division through dilution of the starting signal. Patterns of growth were tracked in continued growth in methanol (left column) or during shifts from succinate to methanol (right column) by measuring fluorescence of individual cells within the populations using flow cytometry at time points over the course of population growth. The resulting data is displayed as distributions of fluorescence intensity that demonstrate how a population does or does not migrate (grow) over time. During continued growth on methanol, there is a uniform decrease in the fluorescence distribution for the whole population for each of the three strains (left column). In contrast, a distinct shoulder or even a second peak of the distribution representing cells that maintain high fluorescence indicates that not all cells have initiated growth simultaneously, if at all. This second mode of the distribution is seen, at least transiently, for all genotypes but is most pronounced in magnitude and time that it remains a part of the population for the *ΔefgA* strain. Data is plotted as histograms of per-cell fluorescence intensity (measured at the emission wavelength of 525 nm, FITC channel) of cell populations at different time points across growth. Each timepoint in a panel represents cell events surveyed within an equivalent volume of growing culture volume. Each panel depicts a single representativebiological replicate.

**Figure S5. Evolved strains with mutations in other loci leading to enhanced formaldehyde growth are not defective in transition to methylotrophy.**

Strains with single-base pair mutations in *def* (CM3908, brown diamonds) and *efgB* (CM3783, green inverted triangles and CM3837, lime-green triangles) were screened for their influence on lag times during carbon source switch that required the cells to transition to methylotrophic growth.

The wild-type (CM2730, black, no symbols), *ΔttmR* mutant (CM4732, purple squares), and *ΔefgA* mutant (CM3745, blue triangles) were included as controls and/or reference points. Succinate-acclimated strains were subcultured into fresh liquid MP medium with methanol as the carbon source and growth was assayed by absorbance. We used a >15% increase from starting absorbance (dashed line) constituted a threshold that marked the end of lag phase. *Error bars* represent the standard error of mean of biological replicates.

**Table S1. Bacterial strains^a^ and plasmids^b^.**

**Table S2. Growth of the *ΔttmR* mutant is comparable to wild-type in the absence of formaldehyde stress.**

*M. extorquens* was grown in MPIPES medium with 3.5 mM succinate or 15 mM methanol as the sole source of carbon and energy. Error shown represents standard from mean of biological replicates. Error was not provided for lag time due to poor resolution at low cell densities. A two-tailed Student’s t-test was performed to identify statistically significant differences in final yields or growth rates between growth rates (p-value < 0.05).

**Table S3. The *ΔttmR* mutant does not mediate a general stress response.**

The wild-type and the *ttmR^EVO^* mutant were subjected to various stressors, including a panel of antibiotics, hydrogen peroxide, ethanol, and heat. The resulting stress was phenotypically manifested by growth inhibition as indicated by zones of inhibition (cm), changes in viability (CFU/mL), or qualitative decrease in colony size.

**Table S4. Transcriptomic analysis of the methylotrophy genes.**

## Contributions

Jannell V. Bazurto: Conceptualization, funding acquisition, investigation, methodology, visualization, writing - original draft preparation, writing – review & editing

Eric L. Bruger: Conceptualization, funding acquisition, investigation, methodology, visualization, writing - original draft preparation, writing – review & editing

Jessica A. Lee: Conceptualization, investigation, methodology, writing - original draft preparation, writing – review & editing

Leah B. Lambert: Funding acquisition, investigation

Christopher J. Marx: Conceptualization, funding acquisition, project administration, resources, supervision, writing - original draft preparation, writing – review & editing

## Acknowledgements

We thank Juan E. Abrahante of the University of Minnesota Informatics Institute (UMII) for assistance with the pipeline for RNA-Seq data analysis and Siavash Riazi for assistance with modifying R scripts. We thank members of the Marx laboratory and Lon Chubiz for critical reading of this manuscript. The flow cytometry was carried out at the IBEST Optical Imaging Core at the University of Idaho (IBEST is supported in part by NIH COBRE grant P30GM103324). This work was supported by funding from an Army Research Office MURI sub-award to CJM (W911NF-12-1-0390), a CMCI Pilot Grant to CJM (parent NIH award P20GM104420), an INBRE Undergraduate Research Fellowship to LBL (parent NIH award P20GM103408), and a BEACON Center for Evolution in Action Pilot Grants to JVB and ELB (NSF Cooperative Agreement DBI-0939454).

